# Development of a 3D-printed Polymeric Heart Valve Structures: From Material Screening to Valve Prototype Evaluation

**DOI:** 10.64898/2026.11.23.768508

**Authors:** Mehdi Saadi, Raymond Haddad, Nour Zeaiter, Oualid Haddad, Divya Prabhakar Reddy, Jessica Bamba-Funck, Angela Sutton, Catherine Boisson Vidal, Kévin Bassand, Hanna Hlawaty

**Affiliations:** Université Sorbonne Paris Nord, Hypoxie & Poumon, INSERM, U1272, F-93000, Bobigny, France; Université Sorbonne Paris Nord, Laboratory for Vascular Translational Science, LVTS, F-93000, Bobigny, France; Université Paris Cité, F-75018, Paris, France; INSERM, U 1148, F-75018, Paris, France; Institut Universitaire de France (IUF); Filière des Cardiopathies Congénitales Enfants Adultes, Hôpital Marie-Lannelongue, Centre Constitutif du Réseau Maladies Rares Malformations Cardiaques Congénitales Complexes - M3C, Hôpitaux Saint Joseph et Marie Lannelongue, F-92350, Le Plessis-Robinson, France; Faculté de Médecine Kremlin-Bicêtre, Université Paris-Saclay, F-94270, Le Kremlin-Bicêtre, France; Université Paris-Saclay, INSERM, UMRS 1358, Hypertension Pulmonaire: Physiopathologie et Innovation Thérapeutique (HPPIT), Hôpital Marie Lannelongue, F-92350, Le Plessis-Robinson, France; Marie Lannelongue Innovation Center (MALIC), Hôpitaux Saint Joseph et Marie Lannelongue, F-92350, Le Plessis-Robinson, France; AP-HP, HUPSSD, Hôpital Avicenne, Laboratoire de Biochimie, F-93000, Bobigny, France; Université Paris Cité, Optimisation Thérapeutique en Neuropharmacologie, INSERM, UMRS 1144, F-75006, Paris, France

**Keywords:** 3D printing, Polymeric printed valves, Low-molecular-weight fucoidan, VEGF165, Coating, Endothelial colonization, Valved-stent

## Abstract

The development of personalized heart valve prostheses remains challenging due to the need to combine anatomical adaptability, suitable material properties, and biological compatibility. To address these limitations, this study investigated the feasibility of developing three-dimensional (3D)-printed polymeric valve prototypes and their biofunctionalization with low-molecular-weight fucoidan (LMWF) and recombinant human vascular endothelial growth factor 165 (rhVGEF165) to support endothelial colonization.

The aim of this study was to evaluate the feasibility of 3D-printing polymeric valve prototypes, identify suitable materials for endothelial cell culture, and develop an LMWF/rhVEGF165-based coating strategy to enhance endothelial colonization.

A trileaflet valve prototype was fabricated using PolyJet printing, and candidate polymeric materials were screened for their physicochemical characteristics, autofluorescence, stability, and compatibility with human endothelial cells. Silicone 40A resin was selected for subsequent surface modification using alternating LMWF/rhVGEF165 multilayer coatings.

Our results showed that silicone 40A resin supported a homogeneous distribution of spread endothelial cells compared with the other evaluated materials. Dip-coating provided a more uniform coating distribution than drop-loading, while the two-layer LMWF/rhVEGF165 coating promoted endothelial colonization without a clear additional benefit from increasing the number of layers. At 144 h, coatings prepared with 100 µg/mL LMWF supported significantly higher endothelial cell density than uncoated surfaces and coatings prepared with 10 µg/mL LMWF. These responses were associated with differences in F-actin organization and the distribution of VE-cadherin, integrin β1, and syndecan-4, suggesting coating-associated changes in endothelial cytoskeletal, junctional, and adhesion protein organization.

Overall, this study establishes the feasibility of combining additive manufacturing, polymer selection, and LMWF/rhVEGF165 surface modification to support endothelial colonization of 3D-printed materials. This approach provides a promising experimental platform for the development of personalized polymeric heart valve substitutes, although further mechanical, hemodynamic, and biological validation is required.

## 1. Introduction

The development of patient-specific heart valve prostheses requires the integration of anatomical, material, and biological considerations. Three-dimensional (3D) printing offers new opportunities to fabricate complex valve geometries from medical imaging data, potentially enabling the production of prostheses tailored to individual patient anatomy [1–3]. Previous studies have demonstrated the feasibility of additive manufacturing for polymeric heart valve prototypes, highlighting its potential to reproduce complex valvular structures and explore alternative designs [4,5].

However, translating these manufacturing advances into functional cardiovascular implants requires materials that combine appropriate mechanical properties, manufacturing compatibility, and long-term stability. Polymers such as silicone elastomers, polycaprolactone (PCL), and poly(lactide-co-caprolactone) (PLCL) offer distinct mechanical characteristics, degradation profiles, and processing capabilities [6–8]. Beyond their bulk properties, their surface characteristics are particularly relevant because they govern interactions with blood components and surrounding cells.

In particular, the blood-contacting surface should support endothelial cell adhesion, spreading, and organization. Establishing an endothelial layer may contribute to creating a biologically active interface between circulating blood and the underlying material. However, synthetic polymeric surfaces often lack the molecular cues required to promote endothelialization. Surface functionalization with bioactive molecules therefore represents a promising strategy to improve the biological properties of these materials.

Among potential bioactive molecules, low-molecular-weight fucoidan (LMWF), a sulfated polysaccharide derived from brown algae, has attracted interest for cardiovascular applications. Its structural and physicochemical similarities to glycosaminoglycans (GAGs) enable interactions with several heparin-binding proteins, including vascular endothelial growth factor (VEGF). Previous studies have shown that LMWF can modulate endothelial cell migration, proliferation, and angiogenic responses, whereas VEGF plays an important role in endothelial cell survival, proliferation, and migration [9–11]. Their combination may therefore provide complementary biological cues to support endothelial colonization.

The potential of this approach for valvular biomaterials was previously investigated by Marinval *et al.*, who developed fucoidan/VEGF-based polyelectrolyte multilayer coatings on decellularized porcine pulmonary valves. These coatings improved endothelial cell adhesion under flow conditions and cell viability, while exhibiting favorable hemocompatibility-related properties [12]. Nevertheless, whether this functionalization strategy can be successfully applied to synthetic 3D-printed polymers and support endothelial cell colonization remains unknown.

Based on these findings, we hypothesized that the sequential deposition of LMWF and recombinant human VEGF165 (rhVEGF165) could generate a biofunctional interface promoting endothelial colonization of 3D-printed polymeric materials. This study aimed to establish a proof of concept for the fabrication and biofunctionalization of a polymeric aortic valve prototype. First, a trileaflet valve was manufactured, and candidate polymers were screened according to their physicochemical properties, macroscopic stability, intrinsic autofluorescence, and ability to support endothelial cell attachment and spreading. Finally, LMWF/rhVEGF165 multilayer coatings were optimized, and endothelial cell colonization, cytoskeletal organization, and the distribution of junctional and adhesion-associated proteins were investigated on the selected material.

## 2. Materials and methods

### 2.1. Endothelial cell culture

Human umbilical vein endothelial cells (HUVEC (C-12200, PromoCell GmbH, Germany) were cultured in Endothelial Cell Basal Medium 2 (ECBM2; C-22111, PromoCell GmbH, Germany) supplemented with 12 % fetal bovine serum (FBS) and the manufacturer’s supplements. Cells were maintained under standard culture conditions at 37 °C in a humidified atmosphere containing 5 % CO_2_.

### 2.2. Patient-Specific valve design and 3D printing

A patient-specific aortic valve model was reconstructed from medical imaging data using a parametric computer-aided design (CAD) approach in Autodesk Fusion. The digital model incorporated the main geometrical features of the aortic root and valve leaflets, with an aortic annulus diameter of 25 mm, a sinotubular junction diameter of 23 mm, and an annulus height of 15 mm. The aortic wall and leaflet thicknesses were set to 2 and 1 mm, respectively, with an inter-leaflet spacing of 0.2 mm and a leaflet angle of 45°.

The CAD model was subsequently transferred to Bone 3D (Paris, France) for additive manufacturing. A full-scale prototype was fabricated using PolyJet technology on a Stratasys J850 Prime 3D printer (Stratasys, Eden Prairie, MN, United States). Agilus30^TM^, a flexible, rubber-like photopolymer designed for PolyJet printing, was selected as the initial printing material. The resulting prototype was macroscopically inspected from the top, bottom, and lateral views to assess the geometric fidelity of the printed structure relative to the CAD design.

### 2.3. Candidate 3D-printing materials

#### 2.3.1. Material selection and sample fabrication

Following the fabrication of the initial full-scale valve prototype, a range of candidate materials was screened for their suitability in the development of 3D-printed cardiovascular devices. The materials included Agilus30^TM^, BioMed Clear resin, silicone 40A resin, silicone elastomers, poly(lactic acid-co-caprolactone) (PLCL 70/30), poly(lactic-co-glycolic acid) (PLGA), and polycaprolactone (PCL).

Agilus30^TM^ samples were supplied by Bone 3D and Prim 3D, whereas BioMed Clear resin and silicone 40A resin samples were provided by Prim 3D. Silicone elastomers with Shore hardness values of 5A, 10A, and 25A were supplied by Lynxter. PLCL 70/30, PLGA, and PCL samples were provided by Lattice Medical.

For standardized material screening and subsequent biological experiments, square specimens (10 x 10 x 1 mm) were fabricated from each material. Their physicochemical, mechanical, and optical properties, together with their resorbability profiles, were compared using manufacturer-provided specifications. The selection criteria included flexibility, hardness, biocompatibility-related classifications, and degradation characteristics (**Table 1**).

**Table 1.** Physicochemical and mechanical characteristics of candidate materials evaluated for 3D-printed heart valve development. **Sh :** Shore; **σm :** Tensile strength; **εᵦ :** Elongation at break; **E :** Young’s modulus; **T**S : Tear strength; **Tg :** Glass transition temperature; **Tm :**Melting

| Material | Manufacturer / Printing provider | Material class / Composition | 3D-printing process | Mechanical properties | Optical appearance | Biomedical status | Resorbability |
|---|---|---|---|---|---|---|---|
| **Agilus30<sup>TM</sup>** | Stratasys / Bone 3D, Prim 3D | Rubber-like photopolymer | PolyJet | Sh 30-35A, σ<sub>m</sub> 2.4-3.1 MPa, ε<sub>β</sub> 220-270 % | Translucent or coloured depending on formulation | No cardiovascular implantability claim identified | Non-resorbable |
| **BioMed Clear Resin** | Formlabs / Prim 3D | Medical-grade photopolymer resin | SLA | Sh 78D, E 2.08 GPa, σ<sub>m</sub> 52 MPa, ε<sub>β</sub> 12 % | Transparent | Biocompatible for specified skin and mucosal-contact applications | Non-resorbable |
| **Silicone 40A Resin** | Formlabs / Prim 3D | 100 % silicone elastomer | SLA, Pure Silicone Technology<sup>TM</sup> | Sh 40A, σ<sub>m</sub> 5.5 MPa, ε<sub>β</sub> 230%, Ts 12 kN/m | Dark (samples used in this study) | Biocompatible for skin-contact applications | Non-resorbable |
| **Silicone shore 5A, 10A and 25A** | Lynxter | Silicone elastomer | Silicone additive manufacturing | Sh 5A, very soft and highly flexible | Translucent, formulation-dependent | Exact biomedical status depends on the commercial formulation used | Non-resorbable |
| **PLCL 70/30** | Lattice Medical | Poly(L-lactide-co-ε-caprolactone) | FDM / filament deposition | Flexible, E 0.4 GPa, σ<sub>m</sub> 15 MPa, ε<sub>β</sub> 1100 %, Tg 30-40°C, Tm 150-170°C | Semi-transparent, slightly white | Medical-grade | 12 - 24 months |
| **PLGA** | Lattice Medical | Poly(lactic-co-glycolic acid) | FDM / filament deposition | Rigid, E 1.5 GPa, σ<sub>m</sub> 55 MPa, ε<sub>β</sub> 6 %, Tg 60–65°C, Tm 150–160°C | Semi-transparent, slightly white | Medical-grade polymer; supplied material not intended for human implantation | 12 - 24 months |
| **PCL** | Lattice Medical | Poly(ε-caprolactone), semi-crystalline thermoplastic | FDM / filament deposition | Flexible, E 0.55 GPa, σ<sub>m</sub> 20 MPa, ε<sub>β</sub> 180 %, Tg < −50°C, Tm 50–70°C | White | Medical-grade | > 36 months, slow degradation |

#### 2.3.2. Intrinsic autofluorescence characterization

The intrinsic autofluorescence of the 3D-printed materials was assessed prior to fluorescence-based biological analyses. Uncoated specimens were imaged using a ZOE^TM^ Fluorescent Cell Imager (Bio-Rad, United States) in the blue (Ex/Em: 380/440 nm), green (Ex/Em: 485/520 nm), and red (Ex/Em: 560/620 nm) channels.

Acquisition settings were kept constant across all specimens within each channel to enable qualitative comparison of intrinsic fluorescence between materials. Phase-contrast microscopy (PCM) was additionally performed to examine surface morphology and printing-related features.

#### 2.3.3. Preliminary evaluation of material stability

The macroscopic stability of candidate materials was initially assessed following 24 h of immersion in ultrapure water. Changes in transparency, color, and structural integrity were visually inspected. Based on this preliminary screening, Agilus30™ specimens were further evaluated after immersion in ultrapure water, phosphate-buffered saline (PBS), endothelial cell basal medium-2 (ECBM-2) without fetal bovine serum (FBS), or ECBM-2 supplemented with FBS. Their macroscopic appearance was monitored at 0, 24, and 48 h. After 48 h, the incubation media were also examined by phase-contrast microscopy (PCM) to detect and characterize the morphology of any material-derived fragments.

### 2.4. LMWF/rhVEGF165 multilayer coating

#### 2.4.1. Coating components

Low-Molecular-Weight Fucoidan (LMWF; 15,500 g/mol) and its 5-(4,6-dichlorotriazin-2-yl)amino fluorescein (DTAF)-labelled form (LMWF-DTAF; 15,500 ± 3,000 g/mol) were kindly provided by Dr. C. Boisson-Vidal (UMRS-1144, Paris, France) [13–15]. Recombinant human vascular endothelial growth factor 165 (rhVEGF165; 293-VE, Bio-Techne/R&D Systems, United States) was used in combination with LMWF to functionalize the 3D-printed samples. The coating procedure was adapted from the polyelectrolyte multilayer deposition method described by Marinval *et al.* [12]. Coating solutions were prepared at final concentrations of 100 µg/mL LMWF and 0.1 µg/mL rhVEGF165. To investigate the effect of fucoidan concentration on endothelial cell colonization, an additional coating condition containing 10 µg/mL LMWF was evaluated while maintaining the rhVEGF165 concentration at 0.1 µg/mL.

#### 2.4.2. Coating deposition methods

Two coating procedures, dip-coating and sequential drop-loading, were compared using 3D-printed silicone 40A resin specimens. For dip-coating, specimens were immersed in 2 mL of LMWF solution (100 µg/mL) for 2 min, followed by rinsing in PBS (pH 7.4) for 2 min. They were then immersed in 2 mL of rhVEGF165 solution (0.1 µg/mL) for 2 min and rinsed again in PBS for 2 min. This sequence was repeated to obtain the desired number of alternating layers.

For sequential drop-loading, 50 µL of LMWF solution (100 µg/mL) was deposited directly onto each specimen and dried for 20 min using a heat gun positioned approximately 30 cm from the surface. The same procedure was subsequently applied using 50 µL of rhVEGF165 solution (0.1 µg/mL). Coatings comprising 2, 6, or 10 alternating layers were prepared using 1, 3, or 5 successive deposition cycles, respectively, each consisting of one LMWF layer followed by one rhVEGF165 layer.

### 2.5. Coating characterization

#### 2.5.1. Microscopy observation

The surface distribution of LMWF-DTAF coatings applied to 3D-printed specimens by dip-coating or sequential drop-loading was examined by fluorescence microscopy using a ZOE^TM^ Fluorescent Cell Imager (Bio-Rad, United States). Surface morphology was characterized by scanning electron microscopy (SEM) using a GeminiSEM 500 microscope (Carl Zeiss, Germany).

#### 2.5.2. Immunofluorescence detection of rhVGEF165

The presence and distribution of rhVEGF165 within the multilayer coatings were assessed by immunofluorescence 48 h after coating preparation. The analysis was performed on 3D-printed Silicone 40A resin specimens.

Coated specimens were incubated with an anti-rhVEGF165 primary antibody (1:50; V4758, Merck, Germany; 60 µL per specimen) for 1 h at room temperature, followed by overnight incubation at 4 °C in a humidified chamber. They were subsequently incubated with Alexa Fluor™ 555-conjugated goat anti-mouse IgG (1:500; A-21424, Thermo Fisher Scientific, United States) for 1 h at room temperature in the dark. Specimens bearing 2, 6, or 10 alternating LMWF/rhVEGF165 layers were imaged using a ZOE^TM^ Fluorescent Cell Imager (Bio-Rad, United States). Parallel specimens incubated with isotype-matched control antibodies were used as negative controls.

### 2.6. Endothelial cell seeding and colonization on 3D-printed materials

#### 2.6.1. Optimization of endothelial cell seeding

Before cell seeding, 3D-printed specimens were rinsed with ethanol, followed by PBS, and individually placed in 12-or 24-well culture plates, depending on the experimental setup. Two cell-seeding approaches were evaluated (Fig. 5A and supplementary Fig. 5). For direct seeding, a 50 µL cell suspension containing 1.5 x10^5^ HUVECs was deposited directly onto the specimen surface.

For insert-assisted seeding, a silicone insert comprising two compartments was positioned on the specimen surface to confine the initial cell suspension. A total of 1.5 x10^5^ HUVECs was seeded per specimen, with 25 µL of cell suspension deposited into each compartment. Cells were allowed to adhere for 4 h at 37 °C in a humidified atmosphere containing 5% CO_2_. Subsequently, 1 mL of complete culture medium was added to each well.

#### 2.6.2. Cell labeling and endothelial colonization

To assess endothelial cell attachment and colonization, HUVECs were labeled with CellTracker^TM^ Green CMFDA (C7025, Thermo Fisher Scientific, United States, 1:2000 dilution) prior to seeding. For the initial material screening, labeled HUVECs were cultured on uncoated Agilus30^TM^, silicone 40A resin, silicone elastomer, PCL, and PLCL 70/30 specimens. Cell distribution and morphology were assessed after 24 h to compare the ability of each material to support endothelial cell attachment. Following material selection, HUVECs were cultured on uncoated and LMWF/rhVEGF165-coated silicone 40A resin specimens using the previously established seeding conditions.

HUVEC colonization was monitored by fluorescence microscopy (FM) at days 1, 4, and 10 during multilayer-number optimization experiments, and at 24, 48, 72, and 144 h during experiments investigating the effect of LMWF concentration. Cell density was determined by counting CellTracker^TM^ Green-labeled HUVECs in fluorescence images and normalizing the cell number to the analyzed surface area. Results were expressed as cells/mm².

### 2.7. Endothelial cytoskeletal and adhesion-associated protein organization

#### 2.7.1. F-actin staining

Forty-eight hours after cell seeding, HUVECs cultured on uncoated and LMWF/rhVEGF165-coated silicone 40A resin specimens were fixed with 4% paraformaldehyde for 20 min at 4 °C and subsequently washed with PBS. Following cell permeabilization, filamentous actin (F-actin) was stained with Alexa Fluor^TM^ 568-conjugated phalloidin (1:400; A12380, Thermo Fisher Scientific, United States) for 20 min at 37 °C in the dark. Cell nuclei were then counterstained with DAPI (10 µg/mL) for 15 min at 37 °C in the dark.

#### 2.7.2. Immunofluorescence

The distribution of proteins involved in endothelial cell-cell and cell-matrix interactions was assessed by immunofluorescence 48 h after cell seeding. Cell-cell junctions were investigated by immunostaining for vascular endothelial VE-cadherin (sc-9989), whereas cell-matrix adhesion-associated proteins were examined using antibodies against integrin β1 (sc-374429) and syndecan-4 (SDC-4; sc-12766). All primary antibodies were obtained from Santa Cruz Biotechnology (United States). HUVECs were fixed with 4 % paraformaldehyde (PFA) for 20 min at 4 °C and washed with PBS. Nonspecific antibody binding was blocked by incubation with 0.2 % porcine gelatin for 1 h at room temperature. Cells were then permeabilized with 0.01% Triton X-100 for 5 min.

Samples were incubated overnight at 4 °C with primary antibodies diluted 1:50 in a humidified chamber. Following PBS washes, samples were incubated with Alexa Fluor^TM^ 555-conjugated goat anti-mouse IgG (1:500; Thermo Fisher Scientific, United States) for 1 h at room temperature in the dark. Isotype-matched control antibodies were used as negative controls. Cell nuclei were counterstained with DAPI, as described above. Fluorescence images were acquired to qualitatively assess the cellular distribution of the target proteins.

### 2.8. Statistical analysis

Statistical analyses were performed using GraphPad Prism (version 10; GraphPad Software, United States). Quantitative cell density data were obtained from three independent experiments (n = 3) and are presented as the mean ± standard error of the mean (SEM). Two-way analysis of variance (ANOVA), followed by Tukey’s multiple-comparisons test, was used to evaluate the effects of coating condition and time. A two-sided \**p* value < 0.05 was considered statistically significant.

## 3. Results

### 3.1. Patient-specific design and 3D printing of the aortic valve prototype

A patient-specific aortic valve model was initially developed using computer-aided design (CAD) to define the valve geometry prior to additive manufacturing (Fig. 1A). The digital model incorporated the main anatomical and geometric parameters of the aortic root and valve leaflets, including annulus height (h), aortic annulus diameter (dAA), sinotubular junction diameter (dSTJ), leaflet angle (la), valvular wall thickness (vw), aortic wall thickness (aw), and interleaflet spacing (ls).

**Figure 1.**
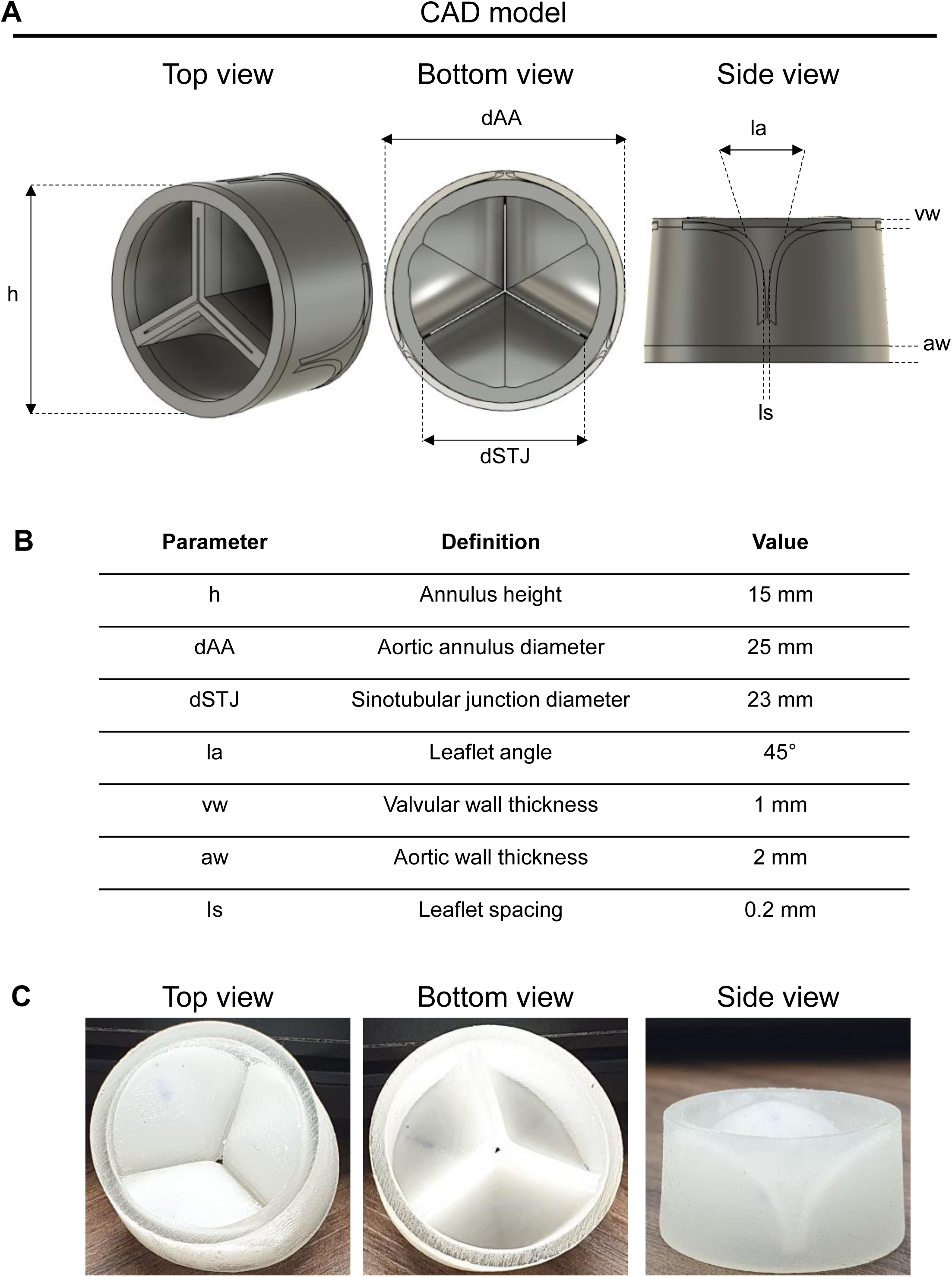
Patient-specific aortic valve design and additive manufacturing. **(A)** Computer-aided design (CAD) model of the aortic valve showing different views and the main geometric parameters defining its parametric design. **(B)** Schematic representation and corresponding values of the geometric parameters used to generate the valve model, including annulus height (h), aortic annulus diameter (dAA), sinotubular junction diameter (dSTJ), leaflet angle (la), valvular wall thickness (vw), aortic wall thickness (aw), and interleaflet spacing (ls). **(C)** Full-scale aortic valve prototype fabricated in Agilus30^TM^ by Bone 3D using PolyJet additive manufacturing technology, shown from different orientations.

The resulting CAD model was characterized by an annulus height of 15 mm, an aortic annulus diameter of 25 mm, and a sinotubular junction diameter of 23 mm. The leaflet angle was defined as 45°, while the valvular and aortic wall thicknesses were set to 1 and 2 mm, respectively. An interleaflet spacing of 0.2 mm was incorporated into the design (Fig. 1B).

The finalized CAD geometry was subsequently fabricated at full scale using PolyJet additive manufacturing technology and Agilus30^TM^ photopolymer material. Macroscopic inspection of the printed prototype from the superior, inferior, and lateral views confirmed the reproduction of the overall valve architecture and the three-leaflet configuration specified in the digital model (Fig. 1C). The individual leaflets and surrounding aortic wall were clearly distinguishable in the printed construct.

Overall, the successful fabrication of the full-scale aortic valve prototype demonstrated the feasibility of translating the patient-specific digital design into a physical construct. This prototype provided the basis for the subsequent assessment and selection of candidate additive manufacturing materials with respect to the structural and biological requirements of aortic valve applications.

### 3.2. Physicochemical properties and autofluorescence screening of candidate 3D-printed materials

Following the fabrication of the initial aortic valve prototype, a panel of candidate materials was selected for the subsequent development of 3D-printed heart valves. Their physicochemical, mechanical, optical, and resorption-related properties were initially compared based on manufacturer-provided specifications (Table 1). The selected materials comprised non-resorbable photopolymers and silicone elastomers, together with biodegradable thermoplastics, including poly(L-lactide-co-ε-caprolactone) (PLCL 70/30), poly(lactic-co-glycolic acid) (PLGA), and polycaprolactone (PCL). These materials encompassed a broad range of mechanical properties, from compliant silicone elastomers (Shore hardness: 5A-40A) to relatively rigid photopolymer resins, such as BioMed Clear resin (Shore hardness: 78D; Young’s modulus: 2.08 GPa). The biodegradable polymers exhibited reported Young’s moduli ranging from 0.4 to 1.5 GPa, with estimated degradation periods ranging from 12-24 months for PLCL and PLGA to more than 36 months for PCL. Their optical characteristics also differed substantially, ranging from transparent and translucent to opaque.

For experimental screening and subsequent cell culture studies, each candidate material was fabricated as standardized square specimens (10 x 10 x 1 mm) to ensure comparable sample dimensions and culture conditions. As subsequent biological investigations relied on fluorescence-based imaging of endothelial cells, the intrinsic autofluorescence of each printed material was characterized by fluorescence microscopy (FM) using blue, green, and red detection channels (Fig. 2). Image acquisition parameters were kept identical within each fluorescence channel to enable direct qualitative comparison of background fluorescence across materials.

**Figure 2.**
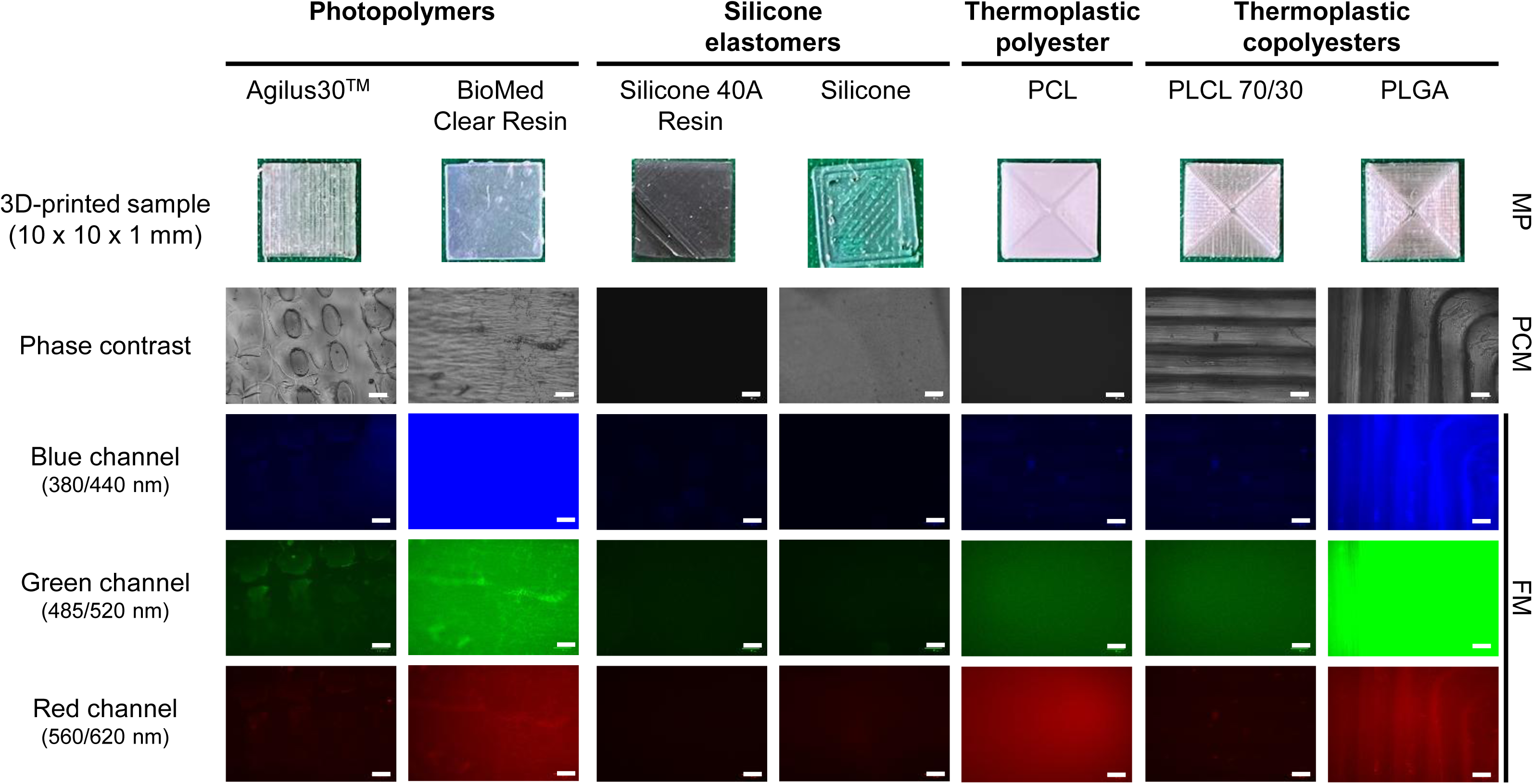
Macroscopic appearance and intrinsic autofluorescence of candidate 3D-printed materials. Macroscopic photographs (MP) of standardized 3D-printed specimens (10 × 10 × 1 mm), classified according to their material families: photopolymers (Agilus30^TM^ and BioMed Clear resin), silicone elastomers (silicone 40A resin and silicone), thermoplastic polyester (PCL), and biodegradable thermoplastic copolymers (PLCL 70/30 and PLGA). Representative images were acquired by phase-contrast microscopy (PCM) and fluorescence microscopy (FM) using blue (Ex/Em: 380/440 nm), green (Ex/Em: 485/520 nm), and red (Ex/Em: 560/620 nm) detection channels with a ZOE^TM^ Fluorescent Cell Imager. Identical acquisition settings were maintained across all materials within each fluorescence channel to enable direct qualitative comparison of their intrinsic autofluorescence. Scale bars: 100 µm.

The fluorescence microscopy analysis revealed distinct autofluorescence profiles among the investigated materials (Fig. 2). For blue channels, BioMed Clear resin and PLGA were autofluroscent. Similarly, PLGA displayed intense fluorescence in both channels, together with a detectable signal in the red channel. Agilus30^TM^ exhibited a heterogeneous fluorescence profile, with a predominant signal in the green channel, whereas PCL showed relatively pronounced diffuse fluorescence in the red channel. By contrast, silicone 40A resin, silicone, and PLCL 70/30 generally exhibited lower background fluorescence, although signal intensity varied across detection channels. Phase-contrast microscopy additionally revealed differences in surface appearance and microstructural features among the printed specimens.

Based on these findings, BioMed Clear resin and PLGA excluded from subsequent fluorescence-based cellular assays because of its pronounced intrinsic autofluorescence, which could interfere with the detection and interpretation of cell-derived fluorescence signals. Also, manufacturer’s indication for PLGA indicates not intended for human implantation. Importantly, the biomedical classifications and material properties reported in Table 1 reflect manufacturer-provided specifications for the corresponding raw materials and should not be interpreted as evidence of regulatory approval or suitability of the resulting 3D-printed constructs for cardiovascular implantation. This initial screening therefore enabled the identification of candidate materials with more suitable optical characteristics and narrowed the material panel for subsequent biological evaluation using standardized 3D-printed specimens.

### 3.3. Preliminary assessment of material stability and HUVEC response on Agilus30^TM^ 3D-printed samples

The macroscopic stability of the remaining candidate 3D-printed materials was initially evaluated following 24 h of immersion in ultrapure water (Fig. 3A). Under these conditions, silicone 40A resin, silicone, PCL, and PLCL 70/30 exhibited no apparent macroscopic changes. In contrast, Agilus30^TM^ showed a visible loss of transparency accompanied by transition into an opaque white appearance, prompting further investigation of its stability under different aqueous conditions.

**Figure 3.**
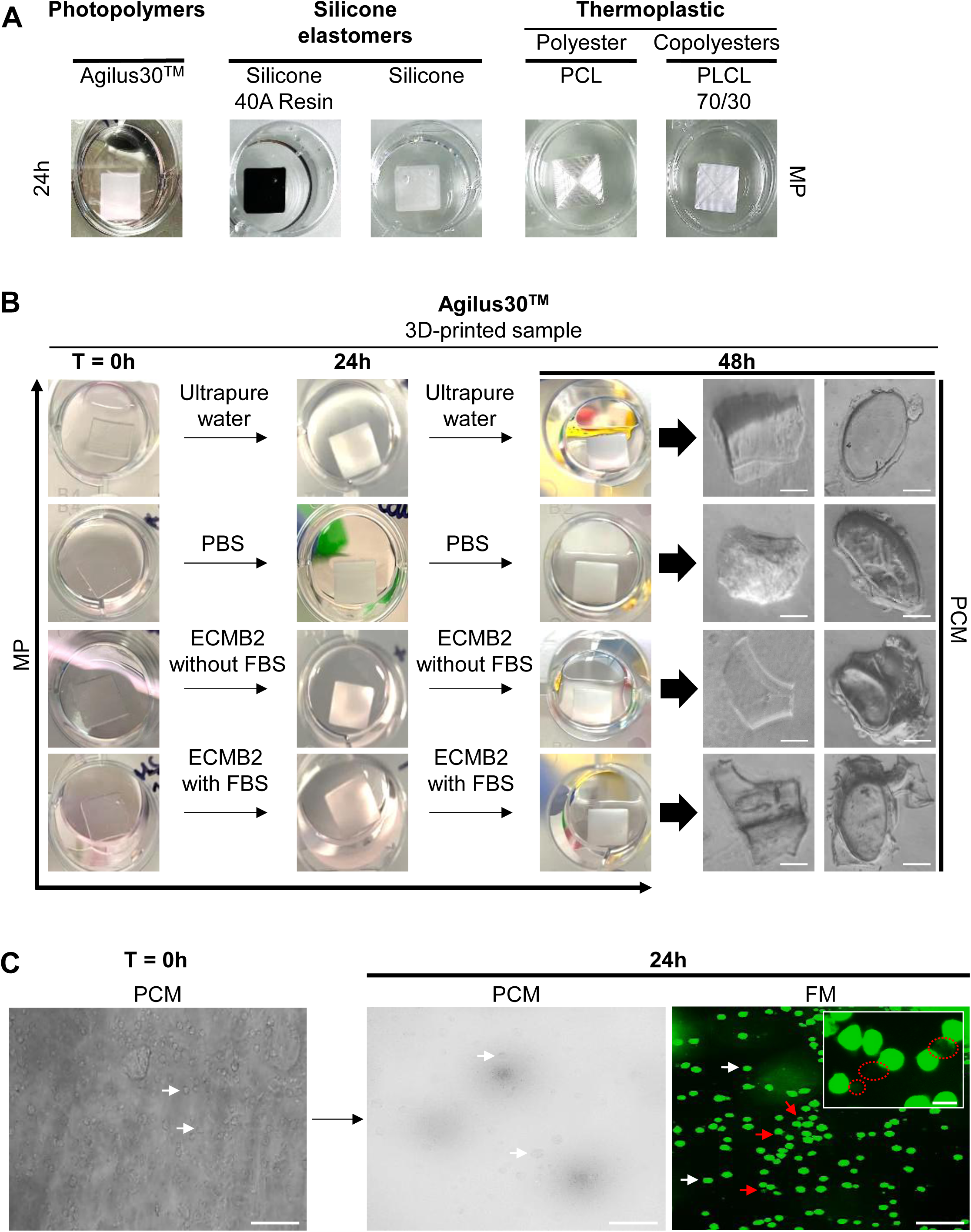
Preliminary assessment of candidate 3D-printed materials and evaluation of Agilus30^TM^ under in vitro conditions. **(A)** Representative macroscopic photographs (MP) of 3D-printed specimens fabricated from different candidate materials following 24 h of immersion in ultrapure water, including the photopolymer Agilus30^TM^, silicone elastomers (silicone 40A resin and silicone), and thermoplastic polyesters, namely poly(ε-caprolactone) (PCL) and poly(L-lactide-co-ε-caprolactone) (PLCL 70/30). **(B)** Representative macroscopic photographs of Agilus30^TM^ specimens at T = 0 h and after 24 and 48 h of immersion in ultrapure water, phosphate-buffered saline (PBS), endothelial cell basal medium 2 (ECBM2) without fetal bovine serum (FBS), or ECBM2 supplemented with FBS. Representative phase-contrast microscopy (PCM) images show material fragments observed in the incubation media after 48 h. Scale bars: 50 µm (left column) and 100 µm (right column). **(C)** Representative phase-contrast microscopy (PCM) images of HUVECs cultured on Agilus30^TM^ specimens at T = 0 h and after 24 h, together with a representative fluorescence microscopy (FM) image of CellTracker^TM^ Green CMFDA-labeled HUVECs after 24 h of culture. White arrows indicate rounded cells exhibiting limited spreading, whereas red arrows indicate small fluorescent structures associated with the cells and displaying altered morphology. The inset shows a magnified view of these structures. Scale bars: 100 µm; inset: 50 µm.

The behavior of Agilus30^TM^ was subsequently investigated following immersion in ultrapure water, PBS, ECBM2 without FBS, and ECBM2 supplemented with FBS for up to 48 h (Fig. 3B). At the initial time point (T = 0 h), the samples appeared predominantly transparent in all four media. However, after 24 h of immersion, visible whitening and loss of transparency were observed under all tested conditions, with these alterations persisting at 48 h.

Phase-contrast microscopy performed after 48 h of immersion further revealed the presence of detached material fragments in the incubation media (Fig. 3B). These fragments exhibited heterogeneous sizes and morphologies, including structures with distinct oval-shaped features. Similar morphological features had previously been observed on the surface of intact Agilus30^TM^ specimens during the initial material characterization (Fig. 2). This morphological correspondence suggests that the detached structures may originate from the printed material. However, their chemical composition and the mechanisms responsible for their detachment could not be established from the available observations.

The interaction of HUVECs with Agilus30^TM^ was subsequently investigated using phase-contrast and fluorescence microscopy following cell labeling with CellTracker^TM^ CMFDA (Fig. 3C). At T = 0 h, cell-like structures were visible on the surface of the printed specimens. Following 24 h of culture, HUVECs predominantly exhibited rounded morphologies, with limited evidence of cell spreading, as observed by PCM.

FM confirmed the presence of CMFDA-labeled HUVECs and revealed additional small fluorescent structures in their vicinity, as indicated by the red arrows and illustrated in the magnified inset (Fig. 3C). Although these features may reflect alterations in cellular morphology or integrity, their nature could not be unequivocally determined from fluorescence imaging alone. Some of these morphological features were reminiscent of changes previously associated with apoptotic or necrotic processes [27]. Nevertheless, in the absence of specific cell-death assays, these observations do not establish the occurrence of cell death or allow discrimination between distinct cell-death mechanisms.

Collectively, the loss of optical transparency and visible whitening of Agilus30^TM^ during aqueous immersion, together with the observation of detached material fragments and limited HUVEC spreading, indicated insufficient material stability and unfavorable initial cell-material interactions under the tested conditions. Accordingly, Agilus30^TM^ was excluded from subsequent biological investigations.

### 3.4. Optimization and characterization of the LMWF/rhVEGF165 multilayer coating

Following the initial material screening, two coating methods, drop-loading and dip-coating, were compared for the deposition of alternating LMWF and rhVEGF165 layers onto 3D-printed silicone 40A resin samples (Fig. 4A).

**Figure 4.**
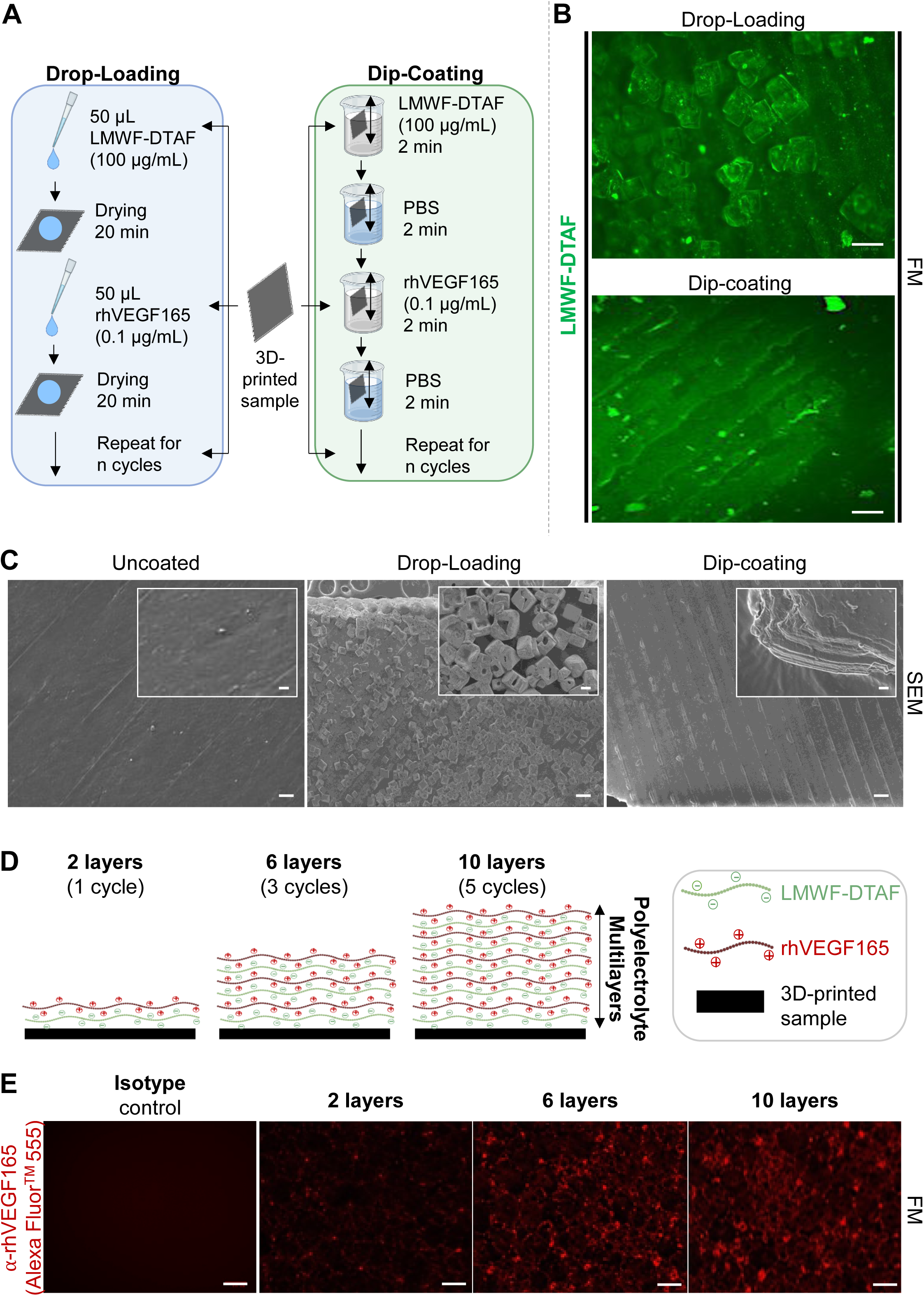
Comparison of coating deposition methods and characterization of LMWF/rhVEGF165 multilayer coatings on 3D-printed silicone 40A samples. **(A)** Schematic representation of the drop-loading and dip-coating methods used for the sequential deposition of alternating LMWF and rhVEGF165 layers. For drop-loading, 50 µL of LMWF-DTAF (100 µg/mL) or rhVEGF165 (0.1 µg/mL) solution was deposited onto the samples, with a 20-min drying interval between successive depositions. For dip-coating, samples were immersed in each solution for 2 min, followed by a 2-min PBS wash between depositions. (B) Representative fluorescence microscopy (FM) images of LMWF-DTAF distribution following drop-loading or dip-coating. Images were acquired using the green fluorescence channel (Ex/Em: 485/520 nm). Scale bars: 100 µm. (C) Representative scanning electron microscopy (SEM) images of uncoated, drop-loaded, and dip-coated Silicone 40A surfaces; higher-magnification views are shown in the insets. Scale bars: 100 µm; insets: 40 µm. (D) Schematic representation of the multilayer coating architecture on 3D-printed Silicone 40A samples. Each deposition cycle consisted of one LMWF layer followed by one rhVEGF165 layer, resulting in coatings comprising 2, 6, or 10 alternating layers after 1, 3, or 5 deposition cycles, respectively. (E) Representative fluorescence microscopy (FM) images of rhVEGF165 immunostaining in coatings comprising 2, 6, or 10 alternating layers. rhVEGF165 was detected using an anti-rhVEGF165 primary antibody followed by an Alexa Fluor^TM^ 555-conjugated secondary antibody. An isotype-matched control was included to assess non-specific staining. Images were acquired using the red fluorescence channel (Ex/Em: 560/620 nm). Scale bars: 100 µm.

The surface distribution of LMWF was first assessed by fluorescence microscopy using DTAF-labelled LMWF (LMWF-DTAF; Fig. 4B). Drop-loading resulted in a heterogeneous fluorescence pattern, characterized by localized accumulations and irregularly distributed fluorescent aggregates. In contrast, dip-coating produced a more continuous fluorescence signal that appeared to follow the surface features of the 3D-printed material, with fewer prominent localized accumulations.

Surface morphology was further investigated by SEM (Fig. 4C). Uncoated Silicone 40A samples exhibited a relatively homogeneous surface, with visible features associated with the 3D-printing process. Drop-loaded samples displayed numerous angular, crystal-like structures, particularly at higher magnification. These structures were markedly less apparent following dip-coating, while the underlying surface morphology remained visible. Overall, these observations suggested a more homogeneous surface distribution of the deposited material after dip-coating, whereas drop-loading appeared to promote localized accumulations. On this basis, dip-coating was selected for subsequent LMWF/rhVEGF165 multilayer deposition.

The selected method was then used to prepare coatings with different numbers of alternating LMWF and rhVEGF165 layers (Fig. 4D). Each deposition cycle consisted of one LMWF layer followed by one rhVEGF165 layer. Accordingly, coatings comprising 2, 6, or 10 layers were prepared using 1, 3, or 5 successive deposition cycles, respectively.

The presence and surface distribution of rhVEGF165 were subsequently assessed by immunofluorescence (Fig. 4E). No detectable fluorescence was observed in the isotype control coated with intermediate layers number (6 layers). In contrast, rhVEGF165-associated fluorescence was detected in all three coating configurations. The signal appeared relatively sparse in the 2-layer coating and more widespread and pronounced in the 6- and 10-layer coatings. This observation suggested that increasing the number of deposition cycles may affect the extent of detectable surface-associated rhVEGF165. However, as fluorescence intensity was not quantified, these observations do not allow a quantitative comparison of rhVEGF165 levels between coating configurations.

Taken together, these results supported the selection of dip-coating for LMWF/rhVEGF165 functionalization of silicone 40A resin samples. The multilayer approach enabled the detection of surface-associated rhVEGF165 in all three coating configurations, with a visually broader fluorescence distribution observed at higher layer numbers.

### 3.5. Optimization of HUVEC seeding and endothelial colonization on 3D-printed materials

A silicone insert-assisted seeding procedure was first evaluated by phase-contrast microscopy (PCM) to establish suitable conditions for endothelial cell culture on 3D-printed square substrates (Fig. 5A). The insert confined the initial cell suspension to the substrate surface, limiting cell dispersion onto the surrounding culture surface. This procedure was subsequently used for all cellular experiments.

**Figure 5.**
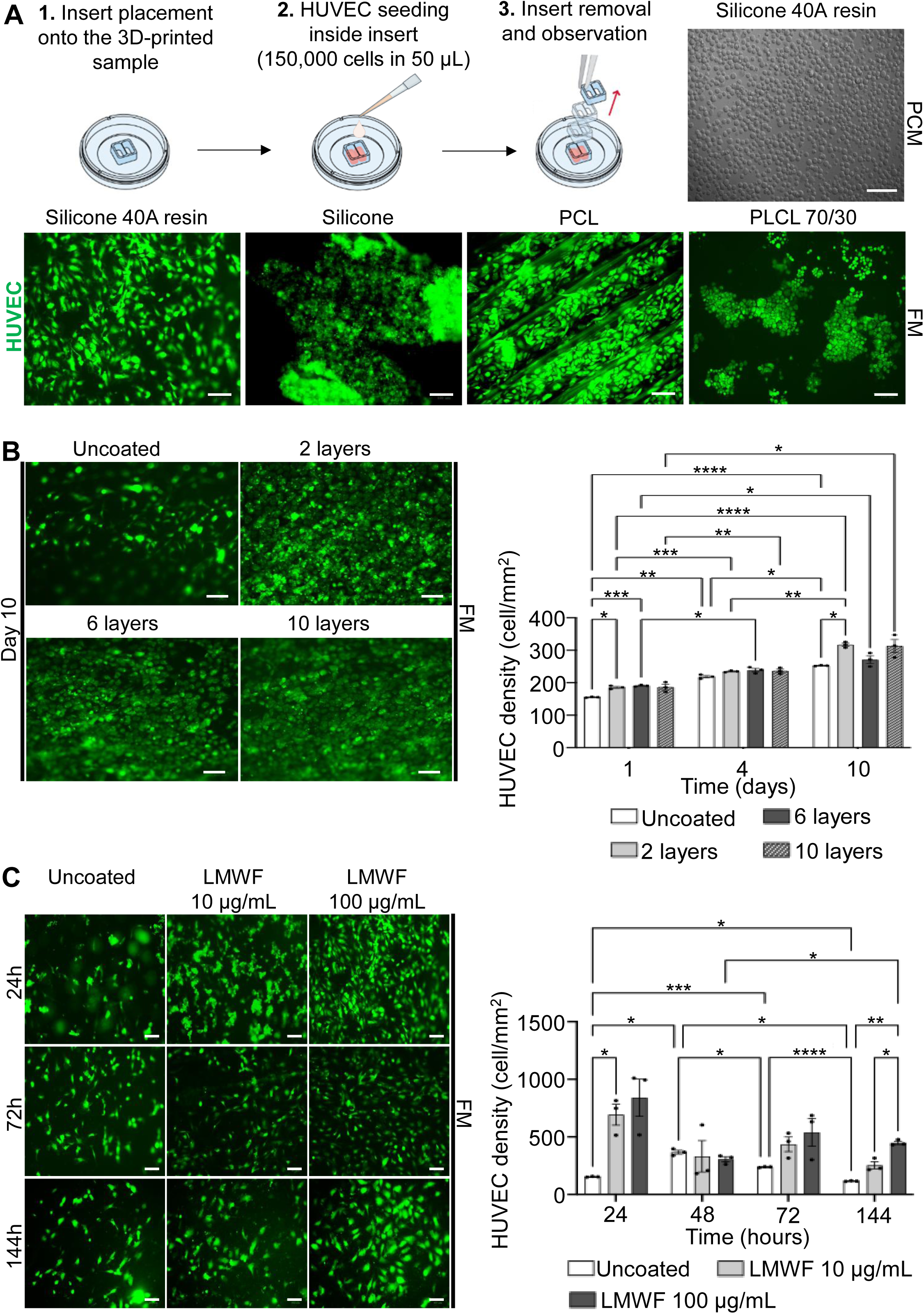
Optimization of HUVEC seeding and assessment of endothelial colonization on 3D-printed candidate materials and LMWF/rhVEGF165-coated silicone 40A substrates. **(A)** Schematic illustration of the silicone insert-assisted HUVEC seeding procedure. HUVECs (150,000 cells in 50 µL) were seeded within an insert positioned on the substrate to confine the cell suspension during initial cell adhesion. The insert was subsequently removed before adding the culture medium. A representative phase-contrast microscopy (PCM) image shows the resulting cell distribution. Representative fluorescence microscopy (FM) images of CellTracker^TM^ Green CMFDA-labeled HUVECs after 24 h of culture on silicone 40A resin, silicone, poly(ε-caprolactone) (PCL), and poly(L-lactide-co-ε-caprolactone) (PLCL 70/30). Pronounced surface depressions and hollow structures on PCL were associated with discontinuous cell coverage. Scale bar: 100 µm. (B) Representative FM images of CellTracker™ Green CMFDA-labeled HUVECs after 10 days of culture on uncoated Silicone 40A substrates and substrates coated with 2, 6, or 10 alternating LMWF/rhVEGF165 layers. Cell density was quantified after 1, 4, and 10 days and expressed as cells/mm². (C) Representative FM images of CellTracker^TM^ Green CMFDA-labeled HUVECs cultured on uncoated silicone 40A substrates or substrates coated with LMWF at concentrations of 10 or 100 µg/mL, with rhVEGF165 maintained at 0.1 µg/mL. Images were acquired after 24, 72, and 144 h, and cell density was quantified after 24, 48, 72, and 144 h and expressed as cells/mm². For panels (B) and (C), data are presented as mean ± SEM from three independent experiments. Statistical significance is indicated as follows: * *p* < 0.05, ** p < 0.01, *** *p* < 0.005, and **** *p* < 0.001. Scale bars: 100 µm.

HUVECs labelled with CellTracker^TM^ Green CMFDA, cultured for 24 h on silicone 40A resin, silicone, PCL, and PLCL 70/30 substrates revealed material-dependent differences in cell distribution and morphology on FM (Fig. 5A). Silicone 40A resin supported the distribution of numerous spread HUVECs across the printed surface. In contrast, silicone and PCL exhibited heterogeneous cell distributions, with localized cell aggregates. PLCL 70/30 also supported HUVEC spreading, with a morphology comparable to that observed on silicone 40A resin. However, pronounced linear topographical features were visible on the PLCL 70/30 surface and coincided with interruptions in cell coverage, potentially limiting the formation of a continuous endothelial monolayer.

Based on these observations, silicone 40A resin was selected for subsequent endothelial experiments, as it supported a more homogeneous distribution of spread HUVECs, without the prominent cell aggregates or surface discontinuities observed on the other materials. This substrate was therefore used for optimization of the LMWF/rhVEGF165 coating.

The effect of the number of alternating LMWF/rhVEGF165 layers on HUVEC colonization was assessed at days 1, 4, and 10 after seeding on Silicone 40A substrates (Fig. 5B). At day 1, HUVEC densities were 186.07 ± 3.37, 191.13 ± 1.68, and 186.91 ± 10.40 cells/mm^2^ on the 2-, 6-, and 10-layer coatings, respectively, compared with 155.69 ± 0.84 cells/mm^2^ on uncoated substrates. The 2-, 6- and 10-layer coatings exhibited significantly higher cell densities than uncoated substrates (\**p* < 0.05 and \*\*\**p* < 0.005, respectively). At day 4, cell densities increased in all groups, reaching 235.44 ± 1.68, 239.24 ± 6.75, and 237.13 ± 6.46 cells/mm^2^ on the 2-, 6-, and 10-layer coatings, respectively, compared with 218.98 ± 4.21 cells/mm^2^ on uncoated substrates. At day 10, the highest mean density was observed on the 2-layer coating (317.72 ± 5.90 cells/mm^2^), compared with 253.16 ± 0.84 cells/mm^2^ on uncoated substrates (\**p* < 0.05), 271.72 ± 14.06 cells/mm^2^ on the 6-layer coating, and 313.92 ± 23.62 cells/mm^2^ on the 10-layer coating. No significant difference was observed between the 2- and 10-layer coatings. Significant increase in cell density over time were also observed.

On uncoated substrates, density increased between day 1 and 4 (\*\**p* < 0.01) and between days 1 and 10 (\*\*\**p* < 0.005). Significant increases between days 1 and 4 were also observed for the 6-layer (\**p* < 0.05) and 10-layer (\*\**p* < 0.01) coatings. Representative FM at day 10 showed extensive cell coverage on coated substrates. Overall, increasing the number of deposited layers beyond two did not provide a clear additional benefit in terms of HUVEC density.

The effect of LMWF concentration on HUVEC colonization was subsequently investigated at 24, 48, 72, and 144 h after seeding, using coatings prepared with two solutions either 10 or 100 µg/mL LMWF and a fixed rhVEGF165 concentration of 0.1 µg/mL (Fig. 5C). At 24 h, HUVEC densities reached 694.51 ± 118.70 and 842.61 ± 216.59 cells/mm^2^ for the 10 and 100 µg/mL LMWF-coated, respectively, compared with 153.58 ± 2.25 cells/mm^2^ on uncoated substrates. A significant difference was reported for the 10 µg/mL condition (\**p* < 0.05). At 48 h, cell densities were 367.93 ± 19.12, 331.64 ± 182.27, and 306.61 ± 29.72 cells/mm^2^ for uncoated, 10 µg/mL, and 100 µg/mL LMWF-coated, respectively. At 72 h, the corresponding densities were 238.81 ± 3.09, 437.13 ± 80.45, and 540.50 ± 159.49 cells/mm^2^. At 144 h, the 100 µg/mL LMWF condition exhibited the highest cell density (449.36 ± 19.40 cells/mm^2^), which was significantly higher than that observed with 10 µg/mL LMWF (253.58 ± 41.91 cells/mm^2^; \**p* < 0.05) and on uncoated substrates (116.87 ± 2.25 cells/mm^2^; \*\**p* < 0.01).

Cell densities varied over the culture period, with both coated conditions exhibiting their highest mean values at 24 h, followed by lower values at 48 h. At the final time point, the 100 µg/mL LMWF condition maintained the highest mean cell density. Representative FM images were consistent with the observed differences in cell coverage. Based on these findings, a 2-layer LMWF/rhVEGF165 coating prepared with 100 µg/mL LMWF was selected for further characterization of endothelial cell morphology and marker distribution.

### 3.6. Junctional, cytoskeletal, and adhesion-associated protein organization in endothelial cells on coated Silicone 40A Resin

Following coating optimization, the organization of the endothelial cytoskeleton, cell-cell junctions, and adhesion-associated proteins was examined by FM on uncoated and two-layer LMWF/rhVEGF165-coated silicone 40A resin substrates (Fig. 6A). F-actin staining was used to assess cytoskeletal organization, while VE-cadherin, integrin β1, and syndecan-4 (SDC-4) immunostaining was used to examine the distribution of junctional and adhesion-associated proteins.

**Figure 6.**
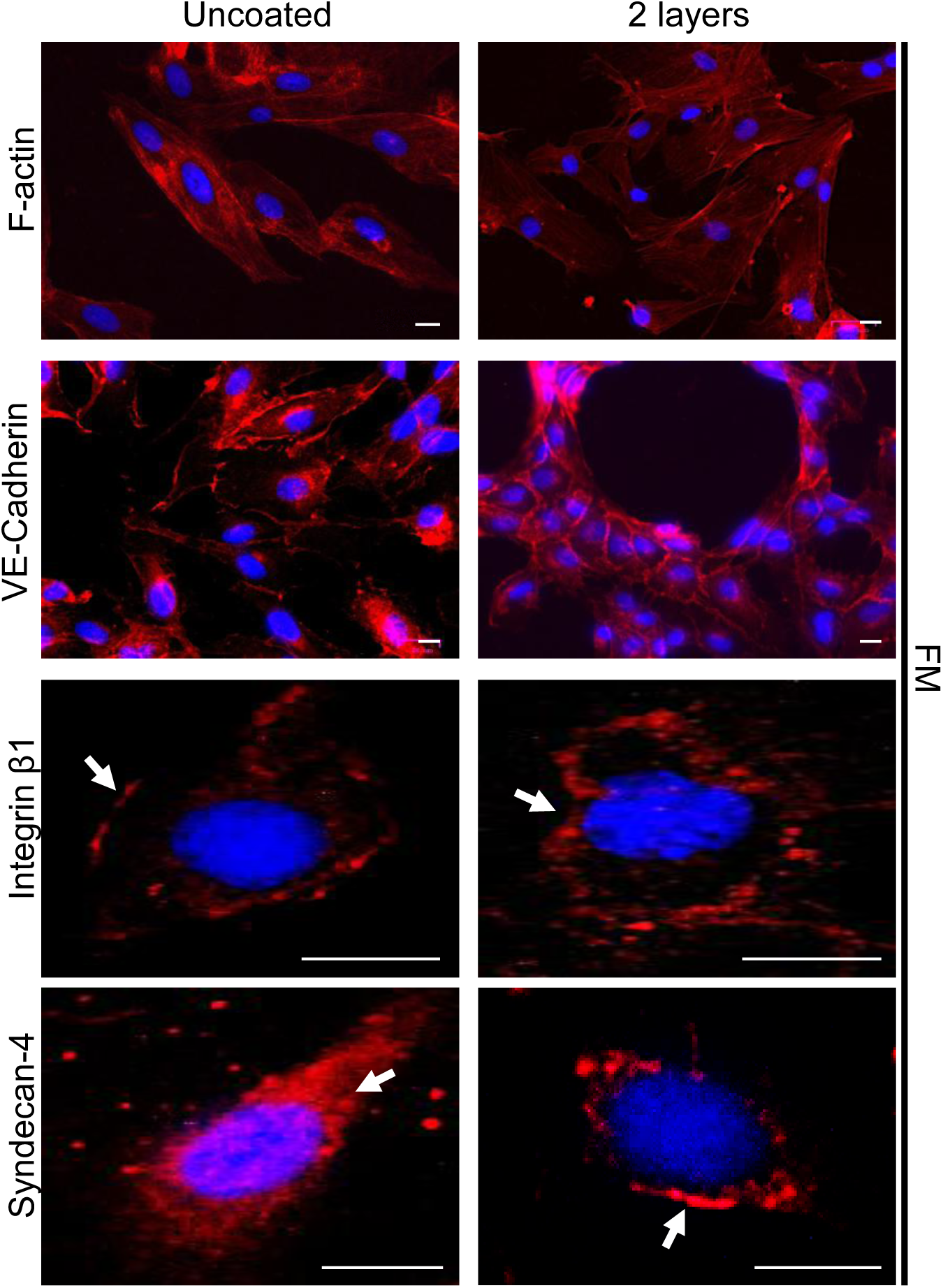
Organization of cytoskeletal, endothelial junctional, and adhesion-associated proteins in HUVECs cultured on coated silicone 40A resin. Representative fluorescence microscopy (FM) images of HUVECs cultured for 48 h on uncoated Silicone 40A Resin or substrates coated with 2 alternating LMWF/rhVEGF165 layers. F-actin, VE-cadherin, integrin β1, and syndecan-4 are shown in red, while nuclei are counterstained with DAPI (blue). Scale bars: 30 µm.

F-actin staining revealed distinct cytoskeletal patterns between the two conditions (Fig. 6A, first row). On uncoated substrates, F-actin was predominantly localized beneath the cell membrane, outlining the cell periphery. In contrast, HUVECs on LMWF/rhVEGF165-coated substrates displayed prominent parallel filamentous bundles extending throughout the whole cell.

VE-cadherin was detected along the cell membrane at cell-cell interfaces in both conditions (Fig. 6A, second row). However, staining appeared visually denser and more intense at cell-cell contacts on LMWF/rhVEGF165-coated substrates. Integrin β1 was predominantly localized at the cell membrane in both conditions, with more pronounced peripheral labeling on LMWF/rhVEGF165-coated substrates (Fig. 6A, third row, white arrows).

SDC-4 staining also differed between conditions (Fig. 6A, fourth row). On uncoated substrates, the signal was predominantly cytoplasmic, with additional labeling overlapping the nuclear region. In contrast, coated substrates exhibited weaker overall staining and a more peripheral distribution, with localized labeling near the plasma membrane (white arrow).

Overall, coated substrates were associated with distinct F-actin organization and qualitative differences in the distribution of endothelial junctional and adhesion-associated markers. These included more prominent parallel actin fibers, visually enhanced VE-cadherin staining at cell-cell contacts, increased peripheral integrin β1 labeling, and a more peripheral SDC-4 distribution. As these observations were based on representative fluorescence images, they do not establish quantitative changes in protein abundance or expression.

## 4. Discussion and conclusion

### ♦ Patient-specific 3D printing as a platform for biofunctionalized aortic valve prototypes

This study establishes an integrated approach combining additive manufacturing, material screening, and biomimetic surface functionalization for cardiovascular device development. A patient-specific trileaflet aortic valve geometry was reproduced by PolyJet printing, while comparative screening identified silicone 40A resin as a promising substrate based on its favorable HUVEC spreading and distribution. LMWF/rhVEGF165 functionalization further supported endothelial colonization, with dip-coating providing more homogeneous apparent coverage and the two-layer configuration showing no consistent disadvantage compared with thicker coatings. Changes in cytoskeletal organization and adhesion- and junction-associated protein distribution suggested additional effects on endothelial organization. In parallel, miniaturized ORMOCER^®^-based stent structures demonstrated the feasibility of fabricating complex supporting architectures. Together, these findings provide a proof of concept for combining 3D printing and surface biofunctionalization in personalized cardiovascular device development, warranting further mechanical and functional validation.

The successful fabrication of the trileaflet prototype demonstrates the ability of additive manufacturing to translate complex CAD-based valve geometries into physical structures. The reproduction of the annulus, leaflets, and surrounding wall highlights the potential of this approach for patient-specific valve development, where anatomical conformity and leaflet geometry are key considerations for subsequent mechanical and hemodynamic performance.

Several additive manufacturing strategies have been explored for polymeric heart valves. Tschorn *et al.* fabricated silicone valve prototypes with different Shore hardness values and evaluated their opening, regurgitation, and transvalvular pressure gradients using a pulse duplicator. Geometric modifications improved hemodynamic performance, although the prototypes did not fully match conventional bioprosthetic valves [4]. Further optimization of the internal architecture subsequently reduced regurgitation and increased the effective orifice area [16].

Additive manufacturing has also enabled indirect fabrication through patient-specific molds. Ilina *et al.* produced silicone models of pediatric heart valves based on three-dimensional ultrasound imaging [17], while Rioux *et al.* used sacrificial printed molds to fabricate patient-specific cellularized hydrogel valves, reproducing anatomical features such as the sinuses of Valsalva while maintaining short-term cellular viability [18]. These approaches illustrate complementary strategies: direct printing enables the fabrication of complex polymeric structures, whereas printed molds facilitate the use of materials that may not be directly printable.

Valve development also requires careful consideration of leaflet mechanics. Cavallo *et al.* assessed the crimping behavior, fatigue resistance, and fluid-dynamic performance of a polymeric aortic valve designed for minimally invasive replacement [5]. Tal *et al.* further demonstrated that biomimetic fiber reinforcement influences strain distribution and valve opening [19]. Together, these studies highlight the need to optimize valve geometry, material properties, and mechanical performance as interdependent design parameters.

### ♦ Physicochemical characterization and material-dependent endothelial responses guide polymer selection

Comparative screening of the 3D-printed materials revealed substantial differences in autofluorescence, aqueous stability, surface morphology, and endothelial responses. BioMed Clear resin and PLGA were excluded because of their pronounced autofluorescence, which limited fluorescence-based characterization. Agilus30^TM^ was also excluded due to its loss of transparency and apparent surface alterations after aqueous exposure, while silicone and PCL showed localized HUVEC aggregates and PLCL 70/30 displayed less homogeneous cell coverage. In contrast, silicone 40A resin provided the most favorable overall appearance and supported the most homogeneous distribution of well-spread HUVECs. It was therefore selected as the primary substrate for subsequent functionalization and endothelial evaluation.

The importance of integrating structural and mechanical characterization in cardiovascular device development has been highlighted by Freystetter *et al.*, who demonstrated that fiber orientation influenced the tensile behavior of 3D-printed and electrospun polyurethane valve scaffolds [20]. Similarly, De Jesus Morales *et al.* developed a biomimetic aortic valve leaflet scaffold combining printed hydrogel structures and PCL [21]. These studies reinforce the need to consider material architecture and mechanical properties alongside biological performance when developing engineered valve substitutes.

Phase-contrast microscopy revealed material-dependent surface features, including linear patterns on PLCL 70/30 and heterogeneous structures on Agilus30^TM^. Although these features may influence cell-material interactions, their dimensions and associated surface roughness were not quantified. Preliminary stability screening further showed that silicone 40A resin, silicone, PCL, and PLCL 70/30 exhibited no apparent macroscopic changes after 24 h, whereas Agilus30^TM^ progressively lost transparency and developed a whitish appearance in ultrapure water, PBS, and endothelial culture media, irrespective of serum supplementation. After 48 h, PCM revealed detached fragments resembling structures observed on intact Agilus30^TM^ surfaces. These alterations may involve swelling, degradation, delamination, or the release of printing-related components, but their origin remains undetermined without further chemical and physicochemical analyses.

The stability of printed polymers and the potential release of residual compounds are important considerations for their biological assessment. Ngan *et al.* reported that optimized post-printing cleaning improved cell viability and reduced the foreign-body response to printed photopolymers [22]. Similarly, Rengarajan *et al.* showed that solvent extraction of leachable compounds reduced cytotoxic effects in human and mouse fibroblast cultures [23]. Although these studies investigated different materials and cell models, they highlight the importance of post-processing in determining cellular responses.

The cellular response observed on Agilus30^TM^ does not establish apoptosis, necrosis, or direct cytotoxicity. Further assessment using viability assays and chemical characterization of released substances would be required to clarify the underlying mechanisms. Nevertheless, its apparent instability combined with limited endothelial spreading supported its exclusion from subsequent investigations.

Among the remaining materials, silicone 40A resin supported the most homogeneous distribution of well-spread HUVECs. By comparison, silicone and PCL showed localized cell aggregates, while PLCL 70/30 supported cell spreading but displayed interruptions in cell coverage coinciding with pronounced linear surface features. These observations suggest that material properties and printing-related surface features may contribute to endothelial colonization, although their respective effects remain to be established.

Previous studies have demonstrated that surface modification and scaffold design can modulate cell–material interactions. Ghaedamini *et al.* reported enhanced hydrophilicity and cell spreading following agarose hydrogel functionalization of 3D-printed PCL scaffolds [24], while Janmohammadi *et al.* showed that pore geometry and alkaline treatment influenced the physicochemical and biological properties of printed PCL [25]. Although these studies did not investigate endothelial cells, they highlight the relevance of surface properties and manufacturing parameters to cellular responses.

Based on its favorable appearance and homogeneous endothelial coverage, silicone 40A resin was selected for subsequent *in vitro* functionalization studies. Its long-term mechanical performance, biocompatibility, and suitability for cardiovascular implantation remain to be established.

### ♦ LMWF/rhVEGF165 multilayer functionalization provides a biomimetic interface for endothelial colonization

Following material selection, alternating LMWF and rhVEGF165 layers were deposited onto silicone 40A resin using two functionalization approaches. FM revealed heterogeneous LMWF-DTAF distribution after drop-loading, with localized accumulations, whereas dip-coating produced a more continuous fluorescent pattern following the printed surface features. SEM also revealed distinct morphologies: numerous angular, crystal-like structures after drop-loading and fewer prominent accumulations after dip-coating. These differences may reflect the influence of deposition procedures on coating redistribution. PBS evaporation during drop-loading could promote local material accumulation, whereas repeated immersion and withdrawal during dip-coating may favor more uniform surface exposure. However, the composition of the angular structures was not determined, and their attribution to crystallized coating components remains hypothetical. Dip-coating was therefore selected based on its more homogeneous apparent coverage, not on demonstrated differences in coating thickness, molecular loading, or retention.

The combination of LMWF and rhVEGF165 was designed to establish a biomimetic interface promoting endothelial responses. LMWF, a sulfated polysaccharide structurally related to glycosaminoglycans, may contribute to growth factor presentation and retention, while VEGF plays a central role in endothelial survival, migration, and vascular remodeling. Their combination may therefore provide a favorable microenvironment for endothelial colonization.

This strategy builds on our previous work, in which we developed fucoidan/VEGF polyelectrolyte multilayers on decellularized porcine pulmonary valves, demonstrating improved HUVEC adhesion under flow, enhanced endothelial viability, antithrombotic properties, and reduced calcification-related responses [12]. The present study extends this approach to a synthetic 3D-printed silicone substrate. Unlike synthetic polymers, decellularized matrices retain extracellular matrix (ECM) components and molecular binding sites that may influence protein retention and cell recognition. Thus, coating formation, biomolecule retention, and biological activity need to be independently evaluated when translating this strategy to synthetic substrates.

The influence of the associated bioactive molecule is further illustrated by Ye *et al.*, who reported that fucoidan/laminin multilayers enhanced endothelial proliferation while reducing platelet adhesion and smooth muscle cell attachment and proliferation [26]. Together, these findings highlight the versatility of fucoidan-based interfaces and suggest that their biological effects depend on both the polysaccharide and the nature of the associated bioactive molecule.

The fluorescence data indicate the presence of rhVEGF165 at the coated surface. Quantitative fluorescence analysis under standardized imaging conditions would allow the rhVEGF165 signal to be compared across coatings comprising 2-, 6-, and 10-layers. To assess coating stability, the retention of both components should be monitored over time. DTAF-labeled LMWF (LMWF-DTAF) could be used to track the fraction of the initial polysaccharide signal remaining at the surface after incubation, alongside measurements of VEGF165 retention. A separate functional assay would help determine whether the retained rhVGEF165 remains biologically active.

Multilayer formation may involve interactions between negatively charged sulfate groups of LMWF and positively charged regions of rhVEGF165, potentially contributing to the association of successive layers [27]. Nevertheless, these interactions were not directly characterized, and other adsorption mechanisms cannot be excluded. Quantitative surface analysis, coating-retention measurements, and rhVEGF165 release assays will be needed to clarify the molecular organization and stability of the interface.

Overall, LMWF/rhVEGF165 multilayers were successfully deposited on a 3D-printed silicone-based substrate, with dip-coating providing more homogeneous apparent coverage. Although rhVEGF165 remained detectable across the tested layer numbers, further quantitative and functional studies are required to determine how multilayer architecture influences biomolecule loading, retention, and endothelial activity.

### ♦ The optimized coating supports time-dependent endothelial colonization

The biological evaluation of functionalized silicone 40A resin first required optimization of endothelial seeding. HUVEC colonization on coatings comprising 2, 6, or 10 alternating LMWF/rhVEGF165 layers showed significantly higher cell densities on 2- and 6-layer coatings at day 1. At day 10, the highest mean density was observed with 2 layers, although it was comparable to the 10-layer condition. Thus, increasing the number of deposition cycles did not proportionally improve endothelial colonization, suggesting that fewer layers may be sufficient to support cell coverage. Additional layers may also alter coating architecture or biomolecule accessibility without necessarily improving their biological availability.

The effect of LMWF concentration was then assessed using 10 or 100 µg/mL, while maintaining a constant rhVEGF165 concentration. Both functionalized conditions showed higher cell densities than uncoated samples at 24 h, with the highest values at 100 µg/mL. Despite a transient decrease at 48 h, this condition maintained the highest density at 144 h, indicating the most favorable colonization profile among those tested. Since cell density reflects attachment, retention, detachment, survival, and proliferation, these changes cannot be attributed to a single cellular process.

These findings are consistent with our previous study of fucoidan/VEGF-functionalized decellularized valvular matrices, which demonstrated improved HUVEC adhesion under flow and enhanced endothelial viability on porcine pulmonary valves [12]. The present study extends this approach to a synthetic 3D-printed silicone substrate under static conditions. Further testing under dynamic flow is needed to determine whether the observed colonization can be maintained under physiologically relevant shear stress.

Overall, the two-layer coating prepared with 100 µg/mL LMWF and rhVEGF165 showed the most favorable endothelial response among the conditions investigated. The absence of a consistent benefit from additional layers suggests that a relatively simple coating architecture may be sufficient to support colonization. However, these findings do not establish coating stability, biomolecule retention, or long-term performance.

### ♦ Coating-associated changes in endothelial cytoskeletal and adhesion-associated protein organization

Beyond differences in endothelial cell density, FM revealed distinct patterns of cytoskeletal organization and adhesion-associated protein distribution in HUVECs cultured on uncoated and LMWF/rhVEGF165-coated silicone 40A resin. These observations suggest that surface functionalization may influence endothelial organization beyond cell colonization.

F-actin staining revealed predominantly peripheral labeling on uncoated samples, whereas cells on functionalized substrates exhibited more prominent filamentous bundles extending throughout the cell body. This difference may reflect changes in cell spreading, cell-substrate interactions, or cytoskeletal organization.

VE-cadherin was detected at intercellular contacts under both conditions, with visually more pronounced labeling on functionalized samples. As a key component of endothelial adherens junctions, VE-cadherin coordinates cell-cell interactions and cytoskeletal organization. Lampugnani *et al.* demonstrated that VE-cadherin regulates endothelial actin organization through Rac activation and Tiam redistribution to the plasma membrane [28]. This provides a mechanistic context for the concomitant differences in F-actin and VE-cadherin staining observed here. Nevertheless, Rac activity and endothelial permeability were not assessed, preventing conclusions regarding junctional function or barrier integrity.

Integrin β1 staining was predominantly localized at the plasma membrane in both groups, with more pronounced peripheral labeling on functionalized samples. Given its role in cell-matrix interactions and adhesion-associated signaling, this distribution may reflect differences in endothelial interactions with the functionalized surface. Further investigation of integrin activation, ligand engagement, and focal adhesion formation, including vinculin and FAK analysis, would help elucidate the underlying adhesion mechanisms

SDC-4 displayed predominantly intracellular staining on uncoated samples, including the nuclear region, whereas functionalized samples showed weaker overall fluorescence and more pronounced peripheral localization. This redistribution may reflect changes in SDC-4 organization at the cell-material interface, potentially involving interactions between LMWF and the endothelial glycocalyx

Fucoidan-based multilayers have been shown to modulate vascular cell responses according to their associated bioactive components. Ye *et al.* reported that fucoidan/laminin coatings enhanced endothelial proliferation while reducing platelet adhesion and smooth muscle cell attachment and proliferation [26]. Unlike laminin, which contains integrin-binding domains, the present coating incorporates rhVEGF165. The observed changes in integrin β1 distribution may therefore arise from distinct interactions, whose respective contributions remain to be elucidated.

Overall, LMWF/rhVEGF165 functionalization was associated with changes in F-actin organization and in the distribution of VE-cadherin, integrin β1, and SDC-4, suggesting modulation of endothelial cytoskeletal and adhesion-associated organization. Whether these changes translate into improved cell–substrate interactions or junctional function remains to be established. Quantitative image analysis, focal adhesion signaling, and endothelial barrier assays will be essential to determine their functional significance.

### ♦ Future perspectives

The present findings support further development of biofunctionalized cardiovascular devices by combining additive manufacturing with LMWF/rhVEGF165 functionalization. Translation to a complete 3D-printed valve will require assessment of coating uniformity on complex geometries and stability under dynamic conditions. Quantification of coating thickness, surface loading, and rhVEGF165 release will also be important to establish relationships between coating architecture and biological performance. Further biological studies should determine whether the observed endothelial colonization leads to a stable and functional endothelial layer. Cell viability, proliferation, barrier integrity, and resistance to physiological shear stress should be investigated, alongside hemocompatibility, inflammatory responses, and calcification-related processes. Single-component controls will help distinguish the respective contributions of LMWF and rhVEGF165. Mechanical and hemodynamic validation will be critical to establish valve performance. Three-dimensional particle tracking velocimetry, as applied by Zeugin *et al.* to a 3D-printed silicone aortic valve [30], could provide insight into flow patterns under pulsatile conditions. Combined with cyclic mechanical testing, this approach would enable assessment of leaflet motion, pressure gradients, regurgitation, and structural durability.

Overall, this study establishes a proof of concept for combining 3D printing and LMWF/rhVEGF165 functionalization to promote endothelial colonization of polymeric substrates. Demonstrating coating stability, endothelial functionality, and mechanical performance under physiologically relevant conditions will be essential to advance this approach towards an integrated valve device.

## Acknowledgements

This work was supported by the Direction de la Recherche et des Études Doctorales (Ministère de l’Enseignement Supérieur et de la Recherche), Université Sorbonne Paris Nord (USPN), and Institut National de la Santé et de la Recherche Médicale (INSERM). Additional financial support was provided by the Institut Universitaire de France (IUF) and the Fondation de l’Avenir. M. Saadi received fellowships from the ministère de l’Enseignement Supérieur et de la Recherche and USPN (SREI 2023). C. Boisson-Vidal’s salary was funded by the Centre National de la Recherche Scientifique (CNRS).

The authors thank Jérémy Adam (Bone 3D), Delphine Prieur (Prim 3D and Lynxter, APHP), Florian Plouviez (Lattice Medical), for their valuable collaboration. We also thank Dr. Michael Trichet (Institut de Biologie Paris Seine, Sorbonne University) for his assistance with electron microscopy, with additional support from the Île-de-France Region and the CNRS. Finally, we sincerely thank Prof. Carole Planes, director of the Hypoxie & Poumon laboratory (INSERM UMRS 1272), for providing access to research facilities.

## CRediT authorship contribution statement

Mehdi SAADI: Writing – original draft, Visualization, Resources, Methodology, Investigation, Formal analysis, Data curation. Raymond HADDAD: Validation, Investigation, Formal analysis, Roles/Writing – original draft. Nour ZEAITER: Investigation. Oualid HADDAD: Investigation, Methodology, Roles/Writing – original draft, Data curation. Divya PRABHAKAR REDDY: Investigation, Roles/Writing – original draft, Software, Data curation. Jessica BAMBA-FUNCK: Formal analysis, Investigation. Angela SUTTON: Resources, Methodology, Roles/Writing – original draft. Kévin BASSAND : Investigation, Methodology, Validation, Data curation, Roles/Writing – original draft, Writing – review & editing. Catherine BOISSON VIDAL : Methodology, Validation, Roles/Writing – original draft, Writing – review & editing. Hanna HLAWATY: Conceptualization, Roles/Writing – original draft, Writing – review & editing, Supervision, Validation, Project administration, Investigation, Funding acquisition, Formal analysis.

---

**Supplementary figure 5. Schematic representation of the direct HUVEC seeding procedure on 3D-printed samples.** The direct seeding method consisted of placing 3D-printed samples in culture wells, followed by the deposition of 50 µL of cell suspension containing 1.5 x10^5^ HUVECs directly onto the sample surface (left panel). Cell distribution was subsequently examined by phase-contrast microscopy (PCM) (right panel). Representative PCM image of HUVEs seeding on 3D-printed samples at T = 0 h. Scale bar: 100 µm.

