## Supplementary figures and images for "Development of a 3D-printed Polymeric Heart Valve Structures: From Material Screening to Valve Prototype Evaluation"

### Supplemental Figure 5

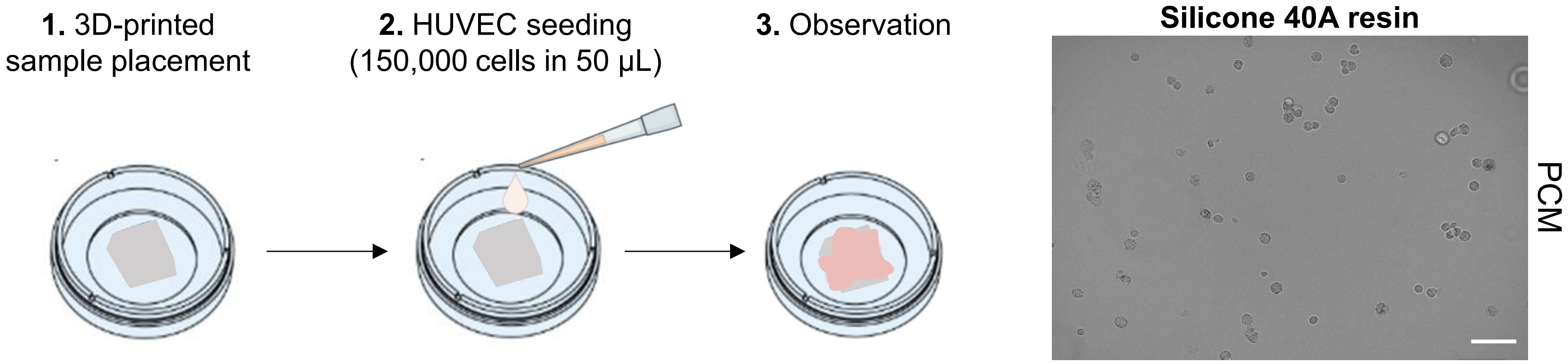
1. 3D-printed sample placement
2. HUVEC seeding (150,000 cells in 50 µL)
3. Observation
Silicone 40A resin
PCM
